# Longitudinal Changes in Nasal and Oral Microbiome and Antimicrobial Resistance Gene Profiles in Response to Human Fecal Microbiota Transplantation

**DOI:** 10.64898/2026.01.27.701854

**Authors:** Mary L. Vallecillo-Zuniga, Ali Akeefe, D. Garrett Brown, Taylor A. Wahlig, Marco Marchetti, Tatum Heiner, Kilee L. Davis, Carmen Nieznanski, Ann Flynn, Daniel T. Leung

**Affiliations:** Department of Internal Medicine, Division of Infectious Disease, University of Utah School of Medicine. Salt Lake City, Utah, United States of America; Department of Biomedical Informatics, University of Utah School of Medicine. Salt Lake City, Utah, United States of America; Department of Human Genetics, University of Utah School of Medicine. Salt Lake City, Utah, United States of America; Department of Internal Medicine, Division of Gastroenterology, University of Utah School of Medicine. Salt Lake City, UT. United States of America; Department of Pathology; Division of Microbiology and Immunology, University of Utah School of Medicine. Salt Lake City, Utah, United States of America

**Keywords:** FMT, gut-lung axis, nasal microbiome, oral microbiome, antimicrobial resistance genes, dysbiosis

## Abstract

The gut-lung axis describes interactions between intestinal and respiratory mucosal systems through microbial, metabolic, and immune pathways, but the systemic impact of gut-targeted therapies on upper respiratory tract (URT) communities remains underexplored. We conducted a longitudinal study in adult patients undergoing fecal microbiota transplantation (FMT) for recurrent *Clostridioides difficile* infection (CDI) alongside healthy controls. Fecal, nasal, and oral samples were collected at baseline (Day 0) and on Days 14 and 56 following FMT. Shotgun metagenomic sequencing was performed to quantify microbial diversity, taxonomic composition, and the abundance of antimicrobial resistance genes (ARGs). FMT was associated with increased gut diversity and decreased levels of key intestinal taxa commonly considered pathobionts, including *Klebsiella* spp., *Escherichia* spp., *Shigella* spp., and *Klebsiella pneumoniae*. At the phylum level, fecal *Bacteroidota* increased, while *Mucoromycota* decreased following treatment. Post-FMT nasal microbiome changes included reduced richness and diversity, expansion of *Moraxella*, and decreases in taxa linked with respiratory colonization, including *Staphylococcus aureus* and *Streptococcus pneumoniae*. By Day 56, nasal communities partially recovered toward healthy profiles. Baseline nasal ARG abundance decreased following FMT, particularly among β-lactam, aminoglycoside, and fluoroquinolone resistance genes, and remained comparable to healthy controls by Day 56. In contrast, the oral microbiome and oral resistome remained largely stable, with only minor fluctuations, and no consistent increases in respiratory pathobiont-associated taxa. In summary, FMT was associated with broader effects beyond the gut, including changes in the URT microbial ecology and antimicrobial resistance profiles. Together, these findings are consistent evidence of gut-lung microbial interactions, linking intestinal dynamics with respiratory microbial composition and antimicrobial resistance patterns.

## INTRODUCTION

The gastrointestinal (GI) microbiota plays an essential role in host health, influencing not only local gut function but also systemic immune, metabolic, and neurological processes.[1–3] The gut-lung axis, which connects the GI and respiratory tracts, has recently emerged as a critical pathway in shaping respiratory immune responses and disease susceptibility.[3–5] These mucosal systems share common structural and immunological features and are colonized by diverse microbial communities that interact through immune signaling, microbial metabolites, and systemic circulation.[5] However, the impact of interventions targeting one system, such as the gut, on another, such as the respiratory tract, remains poorly understood.

Experimental and clinical studies support multiple mechanisms through which gut perturbations influence respiratory outcomes.[6] Gut-derived metabolites such as short-chain fatty acids (SCFAs), cytokines, and microbial components can modulate pulmonary immunity and inflammation.[7] Dysbiosis induced by infection, antibiotics, or chronic disease is associated with increased susceptibility to respiratory infections and inflammatory lung disorders.[8–10] In murine models, depletion of anaerobic gut microbes exacerbates the severity of respiratory infections, even in the absence of significant changes in lung microbiota composition, suggesting immune-mediated mechanisms are involved.[11, 12] Similarly, in human populations, particularly children in low- and middle-income countries, diarrheal diseases have been linked to increased risk of pneumonia.[13, 14]

Despite these insights, few studies have examined how intestinal disturbances affect respiratory microbial communities, which serve as an important barrier against inhaled pathogens.[15, 16] The upper respiratory tract (URT) microbiota, especially in the nasal and oropharyngeal niches, plays a key role in modulating susceptibility and outcomes of respiratory infections. The oropharynx acts as an important source of lung microbiota through microaspiration and mucosal dispersion.[15, 17–19] Specific URT taxa may exert protective or pathogenic effects. For example, colonization with *Staphylococcus aureus* has been associated with reduced influenza severity in models.[13, 20]

Fecal microbiota transplantation (FMT) is a highly effective therapy for recurrent *Clostridioides difficile* infection (CDI), with cure rates exceeding 80%, by restoring gut microbial diversity and function, and offers a unique opportunity to probe systemic microbiome effects.[21, 22] Beyond gastrointestinal restoration, FMT has been shown to influence immune regulation, metabolic pathways, and even neurological and vascular systems.[22, 23] Preclinical studies indicate FMT and gut microbiota restoration can protect against bacterial pneumonia, including *Pseudomonas aeruginosa*, through immune reprogramming and enhanced pulmonary host defense.[9, 11] However, whether FMT also induces compositional or ecological shifts at distal mucosal sites such as the URT remains largely unexplored.

In addition to microbial diversity, the gut microbiome serves as a major reservoir of antimicrobial resistance genes (ARGs), collectively known as the resistome.[24–26] Antibiotic exposure, common among patients with recurrent CDI, enriches gut ARGs, increasing the risk of multidrug-resistant infections and horizontal gene transfer.[23, 27] FMT has been proposed as a strategy to reduce the resistome burden by replacing resistant strains with donor-derived commensals. [28–30] While most studies have focused on fecal resistome dynamics, little is known about whether gut-targeted interventions influence ARG profiles at extra-intestinal sites such as the nasal and oral microbiomes, which may act as reservoirs for clinically relevant resistance determinants.[31, 32] Understanding these systemic effects is crucial for assessing the role of FMT in combating antimicrobial resistance, a global health priority.

In this longitudinal study, we investigated whether intestinal interventions such as FMT are associated with changes in the upper respiratory tract (URT) microbiota. Using a longitudinal cohort of FMT recipients and healthy controls, we examined temporal shifts in nasal microbial richness composition following treatment. These findings suggest that gut-targeted therapies may be associated with microbial changes beyond the intestine, including in the respiratory tract. By characterizing URT microbial dynamics following FMT, this study addresses a critical gap in microbiome research and provides a framework for future studies aimed at understanding how intestinal microbial perturbations relate to respiratory microbial ecology.

## RESULTS

### Characteristics of the Study Participants

Human subjects were enrolled in the longitudinal study titled *Profiling Changes in Respiratory Tract Microbiome in Intestinal Disease*, conducted at the University of Utah. The study was approved by the University of Utah Institutional Review Board (IRB #00129810). The cohort included individuals undergoing fecal microbiota transplantation (FMT) for recurrent *Clostridioides difficile* infection (CDI) and healthy controls recruited from the Salt Lake City area. Demographic and clinical characteristics of the participants, as well as lifestyle and dietary factors, are summarized in **Table 1**, **Supplementary Figure 1,** and **Supplementary Table 1**.

**Table 1.** Demographics and Clinical Characteristics of Participants

| Characteristic | FMT Recipients<br>n =14 (56%) | Healthy Control Group<br>n = 11 (44%) |
| --- | --- | --- |
| **Gender** |  |  |
| Female | 8 (57) | 8 (73) |
| Male | 6 (43) | 3 (27) |
| **Age group, years** |  |  |
| Age (Median & IQR) | 46.5 (37-60) | 34 (30-53) |
| **Self-reported co-morbidities <sup>a</sup>** |  |  |
| Asthma | 2 (14) | 1 (9) |
| COPD | 1 (7) | 0 |
| Acid Reflux | 5 (36) | 3 (27) |
| IBD | 2 (14) | 0 |
| IBS | 3 (21) | 0 |
| Diabetes | 1 (7) | 2 (18) |
| Autoimmune Disease | 4 (29) | 0 |

Notes: FMT, Fecal microbiota transplant. Some participants reported more than one comorbidity

### Microbiome Structure Differs Significantly Across Body Sites

To evaluate the impact of FMT across mucosal compartments, we profiled the microbiomes of stool, nasal, and oral samples longitudinally at baseline (pre-FMT, Day 0), post-FMT (Day 14 and Day 56), along with a cohort of healthy controls (**Figure 1A**). DNA was extracted and sequenced using whole-genome shotgun metagenomics, followed by standard bioinformatic analysis. Principal Coordinates Analysis (PCoA) based on Bray-Curtis dissimilarity revealed distinct clustering of microbial communities by body site (**Figure 1B**). Fecal samples exhibited the most compact and distinct clustering, consistent with the high richness and specialized diversity of the gut microbiome. Nasal and oral samples showed partial overlap, reflecting shared ecological features of the upper respiratory tract. A PERMANOVA test confirmed significant differences in community composition across sites (F = 19.73, p < 0.001), indicating that both anatomical location and treatment status contribute to microbiome structure. Density distribution plots of Abundance-based Coverage Estimator (ACE) microbial richness and Pielou’s evenness further highlighted ecological distinctions across sites (**Figure 1C**). These findings show that fecal samples exhibit the highest richness and the lowest evenness, reflecting the gut’s complex, metabolically diverse ecosystem. Nasal samples had the lowest richness but highest evenness, suggesting a simplified microbial community. Oral communities displayed intermediate richness and evenness, consistent with their transitional role between external and internal environments linking the gut and the URT. These site-specific profiles reflect distinct ecological dynamics across mucosal sites. While differences may have implications for host-microbe interactions, further studies are needed to determine their functional or clinical significance.

**Figure 1.**
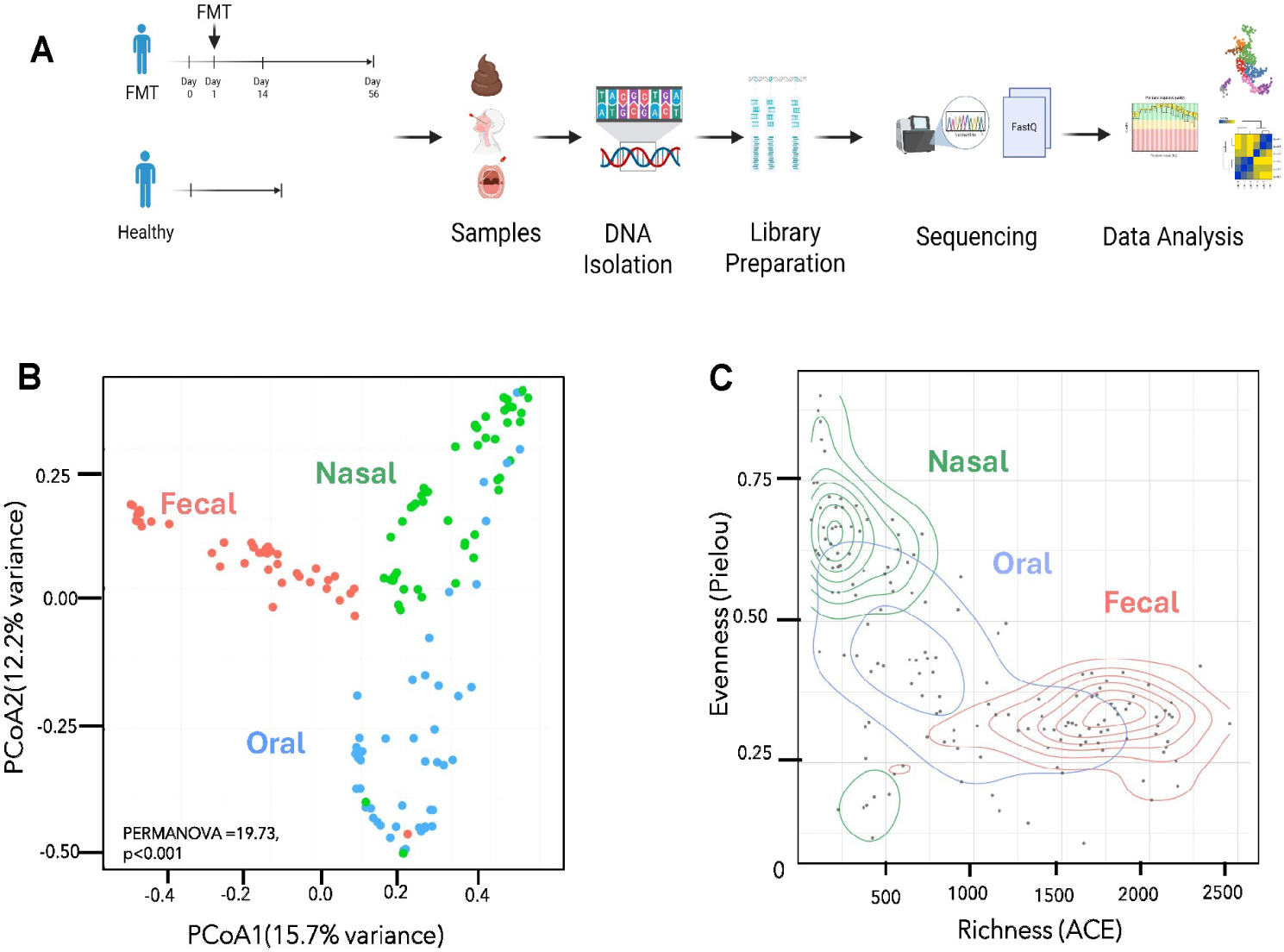
Study Design and Microbiome Overview Across Body Sites. **(A)** Schematic of study design. **(B)** Principal Coordinates Analysis (PCoA) based on Bray-Curtis dissimilarity of microbiome profiles by body site. **(B)** Principal Coordinates Analysis (PCoA) based on Bray-Curtis dissimilarity shows distinct clustering of microbial communities by body site (PERMANOVA F = 19.73, p < 0.001. **(C)** Density distribution plots of microbial richness and evenness across body sites.

### FMT Increases Microbial Richness and Diversity in the Gut while Promoting a Transient Nasal Richness and Diversity Reduction

To assess the systemic effects of FMT on mucosal microbial ecology, we evaluated microbial richness (log₁₀-transformed) and diversity (Shannon Index) across fecal, nasal, and oral samples pre- and post-FMT, as well as in healthy controls. Paired analyses were performed to account for repeated measures within individuals (**Figures 2A and 2B**). Fecal samples exhibited a significant increase in richness post-FMT compared to pre-FMT, approaching levels observed in healthy controls (**Figure 2A**). Fecal Shannon diversity also increased significantly post-FMT (**Figure 2B**), suggesting successful engraftment of donor taxa and improved community evenness, consistent with prior studies demonstrating the restoration of gut microbial complexity and diversity after FMT in patients with recurrent CDI.[33] A notable and relevant finding was observed in the nasal microbiome, where both richness and Shannon diversity decreased by Day 14 post-FMT (**Figures 2A and 2 B**). Although richness and diversity partially rebounded by Day 56, both remained below baseline levels, indicating incomplete recovery within the study period. These results indicate a decrease in nasal richness and a modest downward trend in diversity shortly after FMT. Although partial recovery was observed by Day 56, the extent and duration of this shift remain uncertain. Both measures remained lower than those of healthy controls, suggesting transient changes in respiratory microbial communities following FMT. The oral microbiome exhibited a non-significant reduction in richness, with post-FMT values lower than pre-FMT, while Shannon diversity remained relatively stable, with no significant changes observed (**Figure 2A-2B**). This suggests that the oral cavity is more resistant to ecological perturbations induced by modulation of the gut microbiota. Across all body sites, changes in richness were more prominent than shifts in Shannon diversity. Collectively, these results suggest that while FMT is associated with increased gut microbial richness and diversity, there is a transient reduction in nasal microbial richness, with minimal changes observed in the oral microbiome.

**Figure 2.**
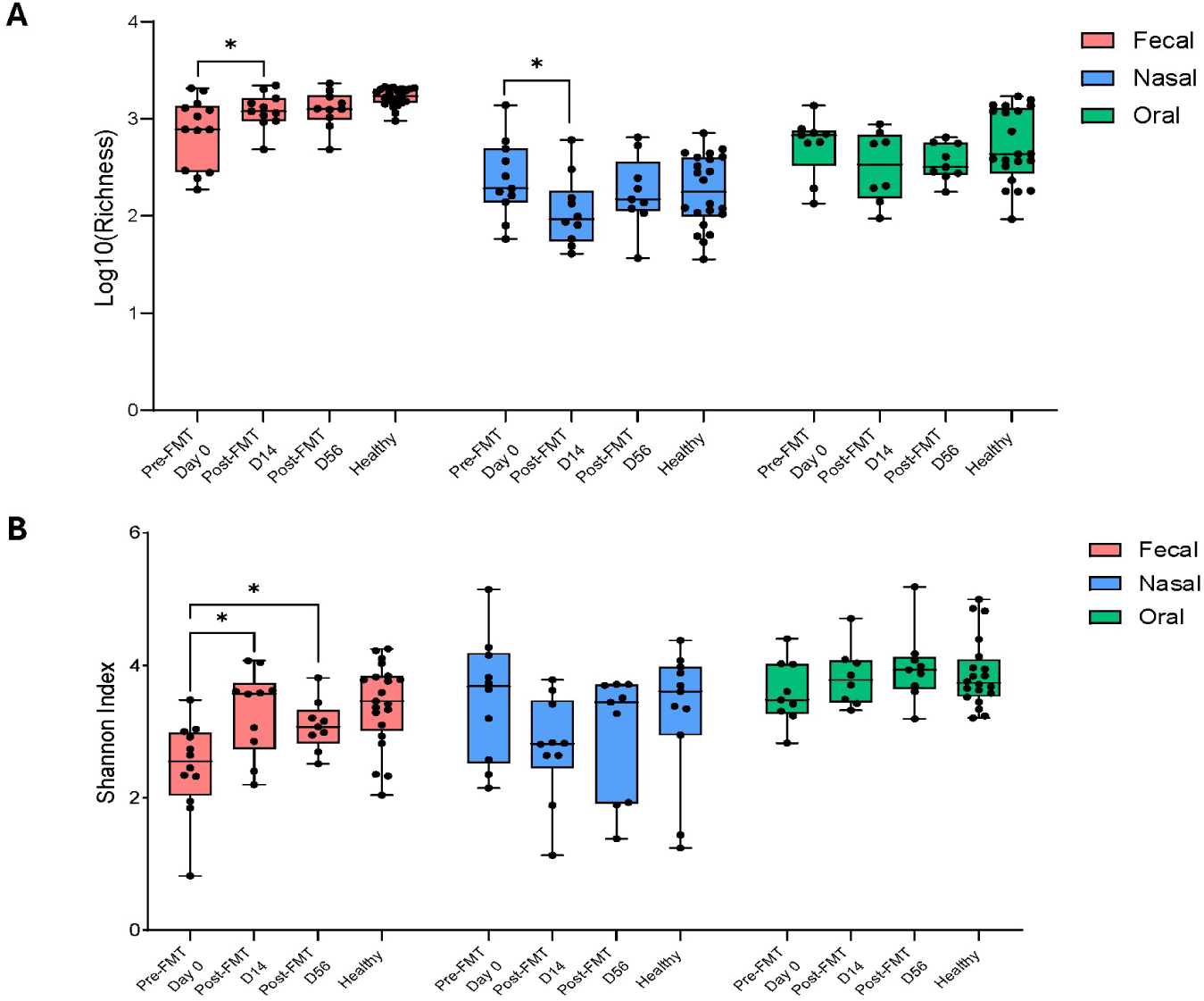
Longitudinal Changes in Nasal and Oral Microbiome Diversity Following FMT. **(A)** Log10 richness and **(B)** Shannon diversity across fecal, nasal, and oral sites at baseline (pre-FMT, Day 0), post-FMT (Day 14 and Day 56), and in healthy controls

### Microbiome Composition and Phylum-Level Shifts Across Body Sites Following FMT

We next evaluated how FMT influenced community composition at the phylum level by calculating the mean relative abundance of bacterial phyla in fecal, nasal, and oral samples pre-and post-FMT and in healthy controls (**Figure 3**).

**Figure 3.**
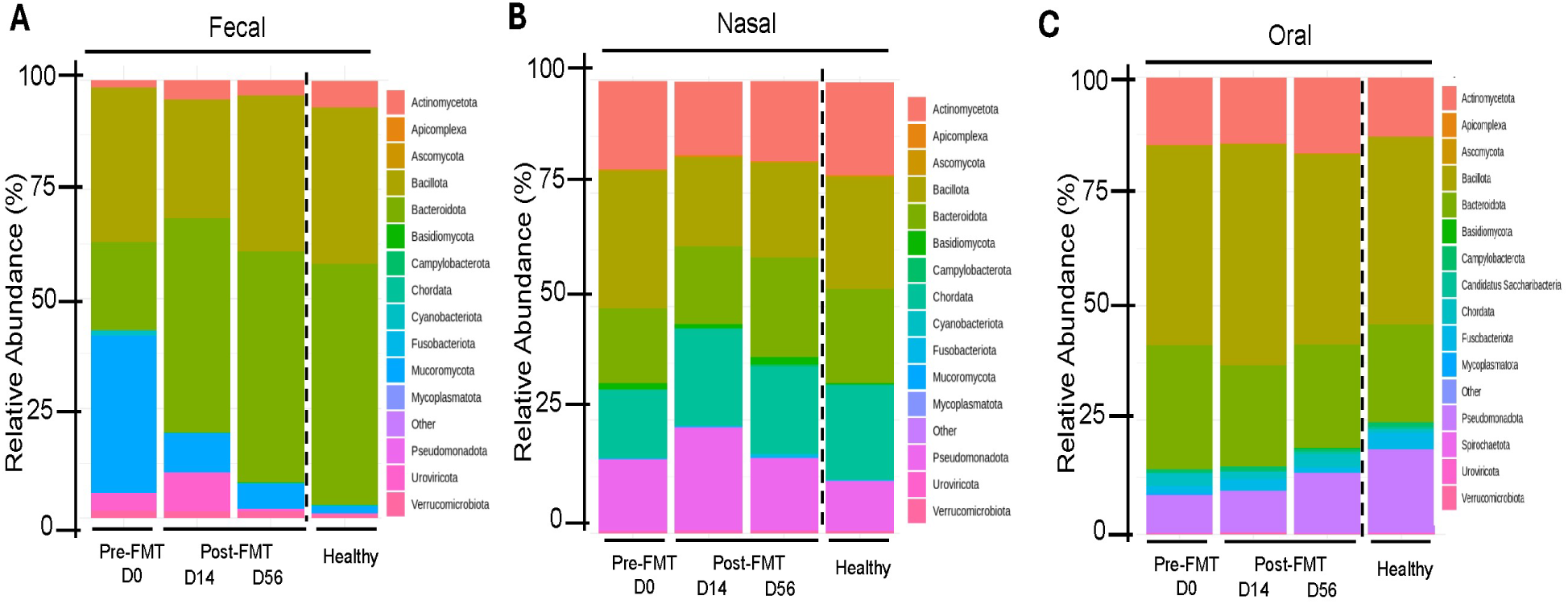
Microbial composition and differential abundance across fecal, nasal, and oral sites following FMT. Mean relative abundance of bacterial phyla in fecal **(A)**, nasal **(B)**, and oral **(C)** samples at pre-FMT (Day 0), post-FMT time points (Day 14 and Day 56), and healthy controls. Each bar represents the average phylum-level composition for the indicated site and time point; colors correspond to phyla listed in the legend. Mean relative abundance at the phylum level across sites and time points (including healthy controls).

The fecal microbiome (**Figure 3A**) was dominated by *Bacillota* pre-FMT, followed by *Bacteroidota*, and *Mucoromycota*, with smaller amounts of *Uroviricota* and *Actinomycetota*. Post-FMT, *Bacteroidota* increased markedly by Day 14 and remained dominant until Day 56, alongside *Bacillota* and *Uroviricota*, while *Mucoromycota* declined sharply. The elevated relative abundance observed at baseline reflects cohort-level means and was influenced by inter-individual variability, rather than uniform dominance across all participants. These changes are consistent with previous reports describing increased relative abundance of *Bacteroidota* and *Bacillota* following FMT in dysbiotic gut microbiomes. [34, 35]

The nasal microbiome (**Figure 3B**) exhibited a distinct phylum-level structure across timepoints, with *Actinomycetota*, *Bacillota*, *Bacteroidota,* and *Pseudomonadota* predominating at baseline. Following FMT, *Bacillota* decreased, while *Pseudomonadota* increased slightly, contrasting with the fecal pattern where *Bacteroidota* expanded and *Mucoromycota* declined markedly (**Figure 3A**). Although nasal microbiome changes were smaller than those observed in the gut, these patterns are consistent with a transient shift followed by partial recovery toward baseline.[36]

Oral samples exhibited a stable phylum-level composition across all time points (**Figure 3C**). *Actinomycetota* and *Bacillota* remained dominant, accompanied by *Bacteroidota* and minor contributions from *Pseudomonadota* and *Fusobacteriota*. Following FMT, the abundance of *Bacteroidota* decreased*. Bacillota* and *Pseudomonadota* increased slightly, making the post-FMT profile more similar to that of healthy controls, which typically display higher *Pseudomonadota* abundance. These results further support the oral cavity’s ecological stability in response to modifications in gut microbiota.

### FMT Reduces Opportunistic Pathogens and Alters Key Genera and Species in the Gut Microbiome

To characterize taxonomic changes following FMT, we examined differential abundance of genera and species in fecal samples before and after treatment (**Figure 4**). Volcano plots (**Figures 4A-B**) revealed significant reductions in opportunistic pathogens, including genera such as *Klebsiella* and *Shigella*, and species such as *Klebsiella pneumoniae* and *Veillonella Parvula*, alongside enrichment of beneficial taxa such as *Phocaeicola* and *Odoribacter*.[37, 38] Boxplot analyses confirmed these findings: *Klebsiella spp.* abundance decreased sharply from pre-FMT to Day 14 and remained low at Day 56 (**Figure 4C**), while *Shigella* exhibited a similar decline (**Figure 4D**). At the species level, *Klebsiella pneumoniae,* a species commonly associated with clinical infection, was significantly reduced post-FMT (**Figure 4E**), and *Veillonella parvula* also decreased (**Figure 4F**). These reductions indicate a decrease in the abundance of several taxa commonly considered potential pathobionts, though the functional relevance of these shifts remains to be determined.[39, 40]

**Figure 4:**
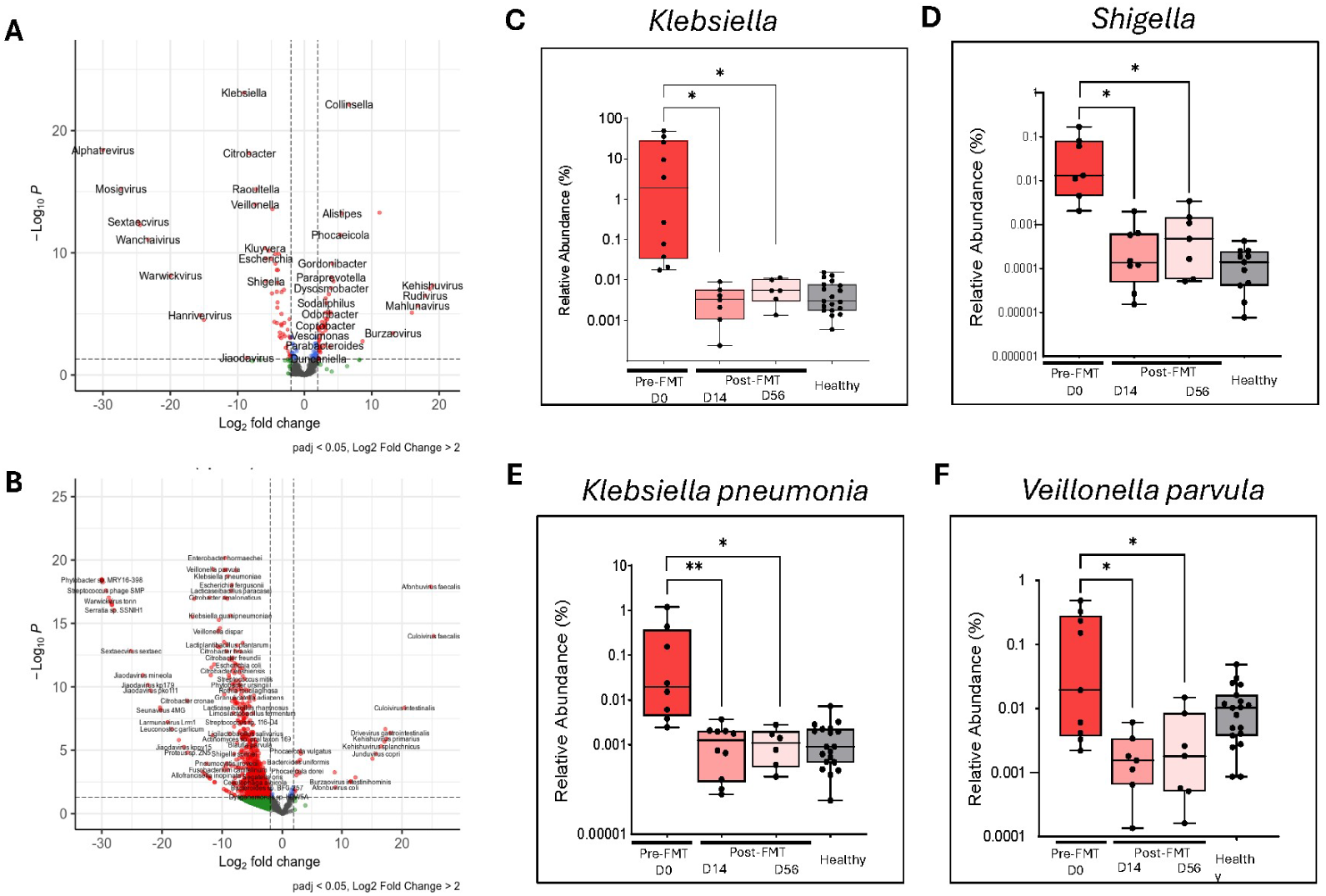
Longitudinal fecal genera and species showing targeted reduction in specific pathogens with FMT. Significant Fecal Genera **(A)** and Species **(B)** comparing Pre- and Post-FMT **(C)** *Klebsiella* spp. **(D)** *Shigella* spp. **(E)** *Klebsiella pneumonia*. **(F)** *Veillonella parvula*

### FMT Induces Selective Shifts in Nasal Microbiome Composition

To evaluate whether gut-target intervention influences the upper respiratory tract microbiome, we examined mean relative abundance patterns of the most dominant nasal taxa at the genus and species levels (**Figure 5A-D**). The top 15 nasal genera, selected on the basis of highest overall relative abundance across all timepoints, demonstrated a stable community structure dominated by *Moraxella, Corynebacterium, Cutibacterium, Bacteroides, Streptococcus*, and *Staphylococcus* (**Figure 5A**). A notable shift occurred post-FMT (Day 14), with *Moraxella* increasing and Staphylococcus and Streptococcus decreasing modestly. By day 56, the nasal community showed partial recovery toward a composition similar to that of healthy controls. We next examined the 10 most abundant respiratory pathobiont genera (**Figure 5B**), which are commonly associated with upper respiratory tract infections, including *Moraxella, Staphylococcus, Streptococcus, Haemophilus,* and *Klebsiella.*[41–48] Consistent with the genus-level trends, *Moraxella* emerged as the dominant pathobiont and further increased after FMT, particularly at Day 14, while *Staphylococcus* and *Streptococcus* decreased. Unlike *Moraxella*, by Day 56, most pathobiont genera trended toward levels observed in healthy subjects, suggesting a transient shift rather than sustained enrichment of these taxa; however, further sampling at later time points post-FMT would be needed to confirm this.

**Figure 5:**
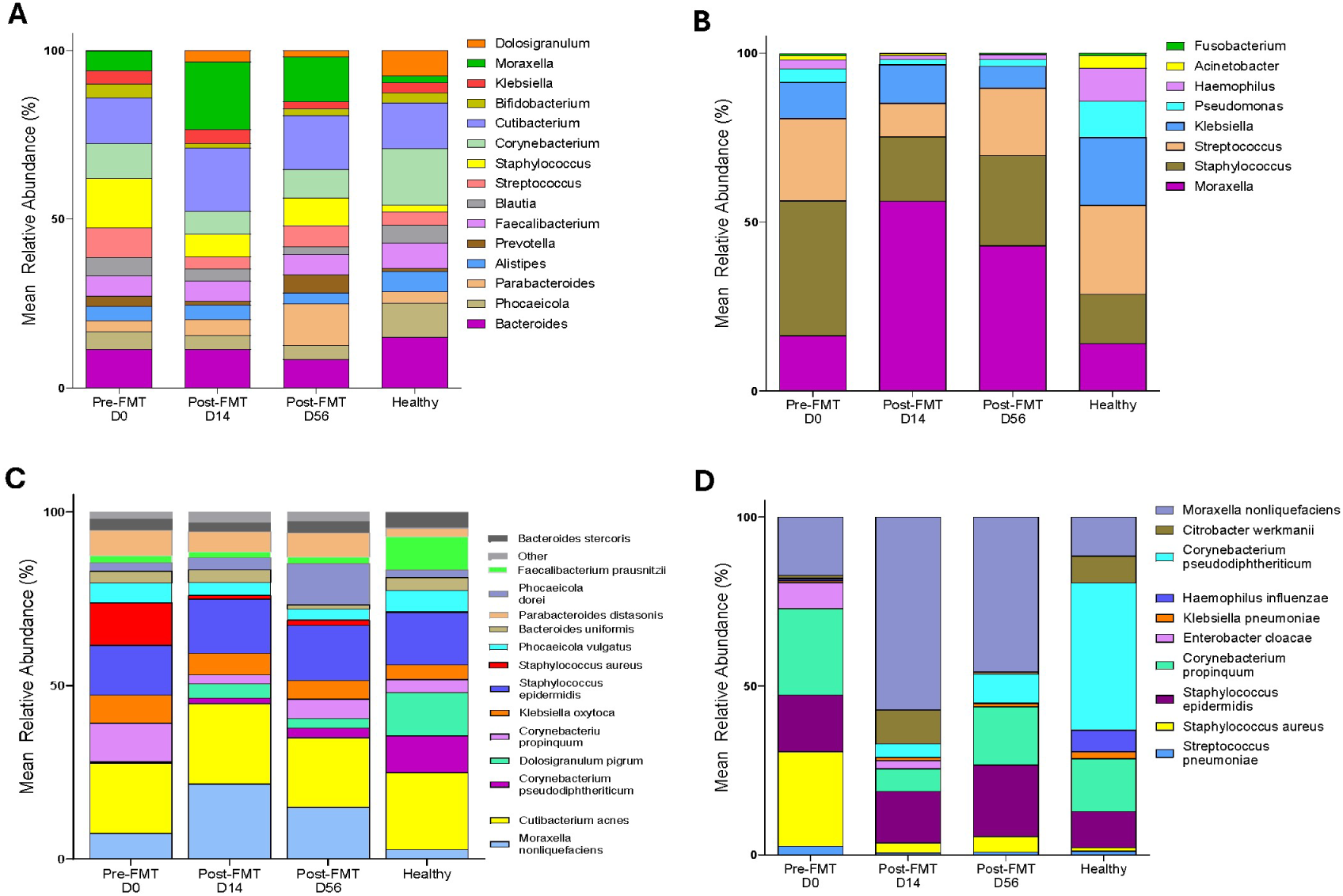
Longitudinal Nasal Mean Relative Abundance. **(A)** Mean relative abundance of the top 15 most abundant nasal genera across Pre-FMT (Day 0), Post-FMT Day 14, Post-FMT Day 56, and Healthy controls. **(B)** Mean relative abundance of the top 10 nasal pathobiont genera, selected from taxa known to include upper respiratory tract opportunistic or pathogenic organisms. **(C)** Mean relative abundance of the top 15 nasal species, designated by the highest overall abundance across all samples. **(D)** Mean relative abundance of the top 10 nasal pathobiont species, including organisms associated with upper respiratory tract infection risk.

At the species level, the 15 most abundant nasal species demonstrated similar patterns (**Figure 5C**). Species such as *Moraxella nonliquefaciens* increased following FMT, whereas *Staphylococcus aureus* decreased, particularly on Day 14 and remained at lower levels at Day 56. These species shifts were not sustained, with partial reversion toward healthy profiles by Day 56. Finally, examination of the top 10 nasal pathobiont species (**Figure 5D**) showed that commensal species, such as *Moraxella nonliquefaciens,* increased, whereas key respiratory pathogens, including *Staphylococcus aureus,* decreased post-FMT. Importantly, no sustained enrichment of clinically significant pathogens was detected on Day 56. Together, these findings indicate that while FMT does not broadly alter nasal microbiome architecture, it is associated with reductions in taxa commonly considered respiratory pathobionts, consistent with cross-site microbial associations following gut-targeted intervention.

### FMT Promotes Limited Changes in Oral Microbiome Composition Without Increasing Respiratory Pathobionts

We next assessed whether FMT altered the composition of the oral microbiome by examining the mean relative abundance of dominant oral taxa at the genus and species levels (**Figure 6A-D**). The top 15 oral genera, selected based on the highest overall relative abundance across all samples, showed a community dominated by Streptococcus, *Prevotella*, and *Rothia,* with minor contributions from *Veillonella, Neisseria, Bacteroides, and Haemophilus* (**Figure 6A**). Although minor compositional fluctuations occurred post-FMT (at Day 14), including a relative increase in *Rothia* and a decrease in *Prevotella,* these changes did not substantially alter the overall community architecture.

**Figure 6:**
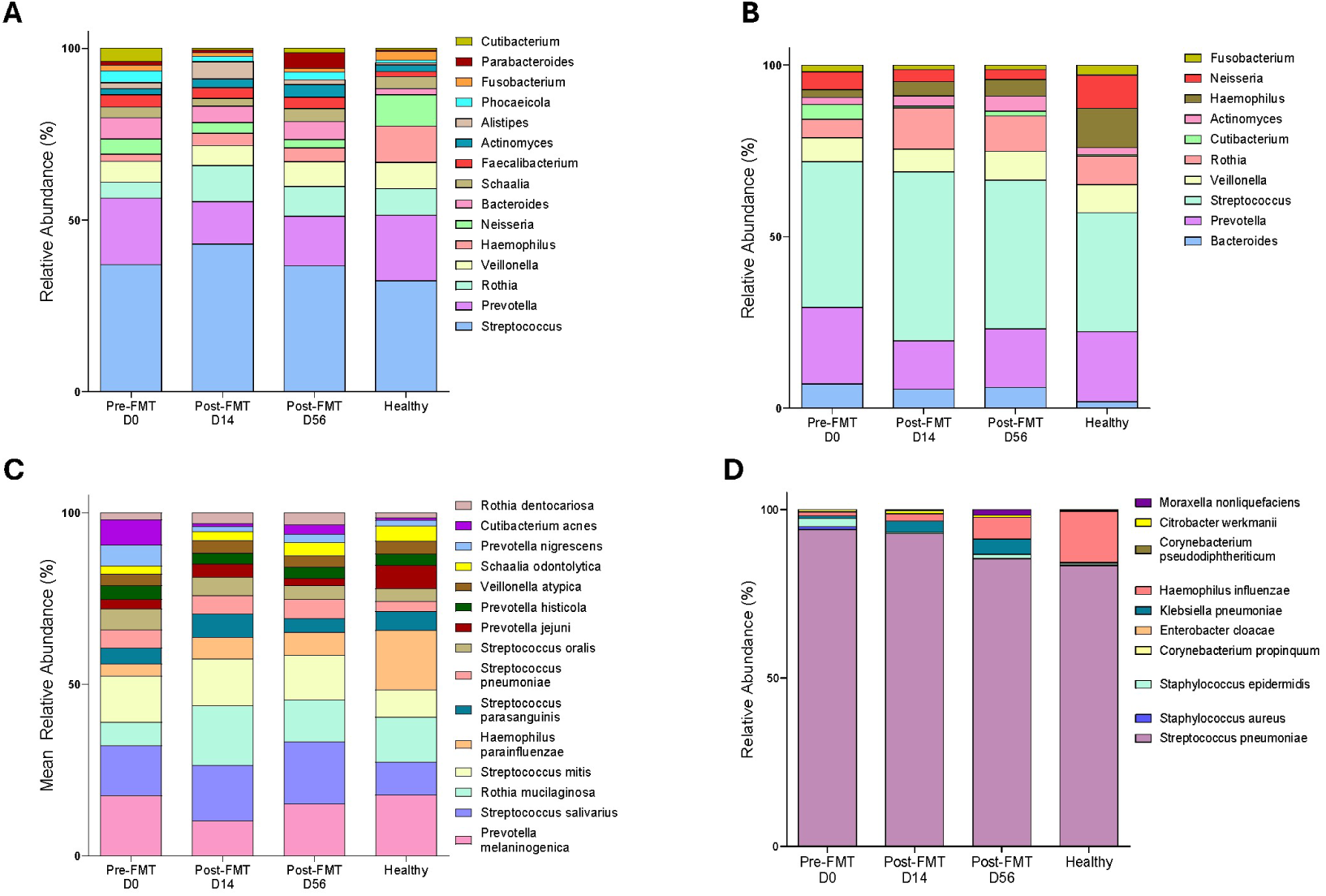
Oral genera and species relative abundance across FMT timepoints. **(A)** Mean relative abundance of the top 15 genera in oral samples across Pre-FMT (Day 0), Post-FMT (Day 14), Post-FMT (Day 56), and Healthy controls. **(B)** Top 10 Pathobionts Oral Genera **(C**) Top 15 Oral Species **(D)** Top 10 Pathobionts Oral Species.

Targeted analysis of the top 10 oral pathobiont genera revealed a similarly stable profile (**Figure 6B**). Genera commonly associated with respiratory or oral opportunistic infections, such as *Streptococcus*, *Prevotella*, *Veillonella, and Haemophilus*,[49, 50] did not exhibit substantial or persistent increases following FMT. However, while *Rothia* and *Prevotella* showed changes on Day 14, most pathobiont genera approached baseline levels similar to those of healthy controls by Day 56.

Species-level profiling of the 15 most abundant oral species (**Figure 6C**) reinforced the observation of oral microbiome resilience. *Rothia mucilaginosa*, a commensal species commonly found in the oral cavity [51, 52], increased noticeably at Day 14 and remained at levels comparable to those of healthy controls. Other dominant species, such as *Prevotella melaninogenica*, *Streptococcus salivarius*, and *Streptococcus mitis*, remained comparatively stable across time points, demonstrating the ecological resilience of the oral microbiome to interventions targeting the gut.

Examination of the top 10 species classified as respiratory pathobionts (**Figure 6D**) was dominated by *Streptococcus pneumonia*, which showed increased relative abundance during the Pre-FMT and Day 14 periods. By Day 56, the levels became comparable to those of the healthy controls. Other pathobionts, such as *Klebsiella pneumoniae* and *Haemophilus influenzae*, remained low before FMT and displayed a modest increase after FMT.

Collectively, these data indicate that the oral microbiome, unlike the nasal microbiome, does not undergo substantial ecological disturbance following FMT. Instead, the oral environment maintains a relatively stable community structure following FMT, with limited and transient changes in dominant taxa, including *Rothia mucilaginosa*. Importantly, these results show no sustained expansion of respiratory pathobionts.

### FMT Reduces Gut Resistome Burden While Nasal and Oral Resistome Remain Low

To understand the systemic impact of FMT on modulating the presence of antimicrobial resistance genes (ARGs) across mucosal sites, we used DNA metagenomic sequencing and the Comprehensive Antibiotic Resistance Database (CARD) to identify and quantify ARG abundance. Total ARG abundance was defined as the sum of log₁₀-transformed ARG abundances normalized per million reads, in fecal, nasal, and oral samples collected longitudinally pre- and post-FMT, and in healthy controls (**Figures 7A and 7C**). As expected, fecal samples showed the highest ARG abundance, possibly reflecting prior antibiotic and healthcare facility exposures in patients with recurrent CDI (**Figure 7A**). Following FMT, total ARG abundance decreased significantly by Day 14 and remained low on Day 56, approaching levels observed in healthy controls. This reduction parallels the decline in taxa such as *Klebsiella* and *Shigella* (**Figure 4**) and is consistent with prior reports of reduced resistome burden following FMT.

**Figure 7:**
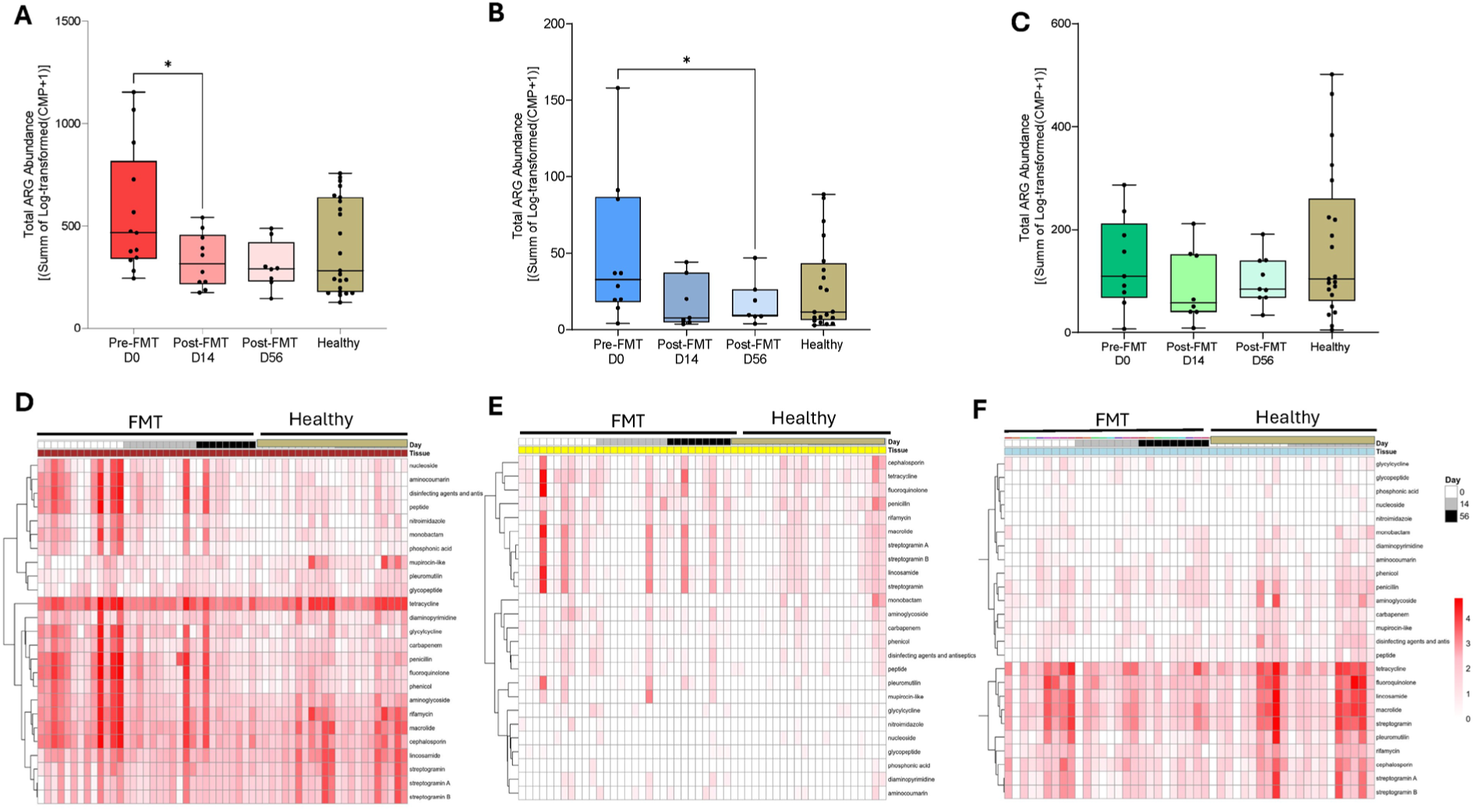
Total ARG abundance in fecal, nasal, and oral samples. (A). Fecal ARG Abundance. **(B).** Nasal ARG Abundance. **(C).** Oral ARG Abundance. **(D-F)** Heatmaps of ARG class-level abundance in fecal, nasal, and oral samples.

Heatmap analysis of fecal ARG class composition (**Figure 7D**) revealed that pre-FMT samples contained high abundances of resistance determinants associated with aminoglycosides, beta-lactams, fluoroquinolones, macrolides, tetracyclines, rifamycins, and glycopeptides, consistent with severe gut dysbiosis in recurrent CDI. Post-FMT samples showed a broad contraction across these classes, forming clusters more similar to those of healthy controls. These findings indicate that FMT is associated with reductions in total ARG burden and contraction of ARG class composition toward profiles observed in healthy controls. [28, 30, 53, 54]

In the nasal microbiome, total ARG abundance decreased by Day 14 post-FMT and was significantly lower at Day 56 (**Figure 7B**), suggesting that gut-targeted interventions, such as FMT, could influence ARG reservoirs beyond the gastrointestinal tract. This pattern mirrors the transient contraction of nasal richness and diversity described earlier (**Figure 2**) and is consistent with changes in upper airway antimicrobial resistance profiles following FMT. Importantly, nasal ARG abundance in post-FMT samples converged toward levels observed in healthy controls. This finding highlights an association between gut-directed intervention and changes in upper airway antimicrobial resistance profiles. Heatmap profiling (**Figure 7E**) revealed that pre-FMT nasal samples were characterized by ARG classes, including beta-lactam, macrolide, tetracycline, and fluoroquinolone resistance genes, indicating a more diverse upper-airway resistome. Following FMT, these classes were moderately reduced over time. Importantly, FMT did not introduce new or clinically concerning resistance classes into the nasal microbiome. These results parallel the reduction in nasal microbial richness (**Figure 2**) and are consistent with coordinated changes in microbial and resistome features across mucosal sites.

The oral ARG abundance exhibited a non-significant downward trend over time. Heatmap analysis further demonstrated that oral samples were dominated by persistent resistance classes, primarily macrolides, tetracyclines, beta-lactams, streptogramins, and aminoglycosides, with minimal change across sampling periods (**Figure 7F**). Together, these findings indicate that the oral resistome remains relatively stable following FMT, in contrast to the more pronounced reductions observed in the gut and nasal microbiomes.

Overall, reductions in ARG burden following FMT were most pronounced in the gut and nasal microbiomes, with more limited changes observed in the oral resistome. This cross-site pattern is consistent with coordinated, site-specific resistome responses across mucosal compartments following gut-directed intervention.

## DISCUSSION

This longitudinal metagenomic study demonstrates that a gut-directed intervention is associated with microbial changes across multiple mucosal sites, including the URT. Although the effects of FMT on fecal microbiome restoration are well established [22, 28, 55], the present study suggests that these gut-level microbial alterations are accompanied by concurrent changes in URT microbial dynamics.

Within the URT, we observed site-specific responses to FMT. The nasal microbiome exhibited a transient contraction in richness and diversity at Day 14 post-FMT, with partial recovery by Day 56. These temporal patterns raise the possibility that gut-targeted interventions may influence distal mucosal ecosystems, potentially through host-mediated pathways, such as immune or metabolic signaling, rather than direct microbial transfer. [4, 7, 9] Taxonomic analysis revealed selective shifts among dominant genera commonly detected in the upper respiratory tract, with *Moraxella* increasing transiently, and *Staphylococcus aureus* decreasing, without sustained enrichment of taxa typically associated with respiratory infections. These patterns align with ecological models of the URT, in which competitive interactions among commensals and pathobionts contribute to colonization resistance.[15, 36, 49] The concurrent reduction in nasal ARG abundance suggests that gut microbial interventions may indirectly modulate upper airway resistome profiles, a finding of interest given the established role of nasal carriage as a reservoir for respiratory pathogens.[31, 32] In contrast, the oral microbiome exhibited marked stability, maintaining core taxa such as *Streptococcus, Prevotella*, and *Rothia* across all time points.[49, 50] Although minor fluctuations were observed in *Rothia mucilaginosa,* a commensal species frequently reported in the oral cavity, [51, 52] overall community architecture and resistome composition remained largely unchanged. This resilience may reflect intrinsic ecological features of the oral niche, including high microbial biomass, frequent nutrient exposure, and strong ecological interactions, which may limit the influence of gut-derived perturbations relative to the nasal environment. [32] Together, these findings support a conceptual extension of the gut-lung axis model beyond immune signaling alone, suggesting that intestinal microbial perturbations may coincide with coordinated microbial and resistome-level changes across mucosal sites. [3, 4, 9] While FMT did not induce large-scale restructuring of URT communities, the observed transient nasal shifts and resistome contraction highlight potential systemic associations that merit further mechanistic and clinical investigation.[28–30]

Importantly, this study has limitations that should be acknowledged. First, the sample size and donor variability were modest and may limit the detection of subtler taxonomic or functional shifts across mucosal sites. Second, the follow-up period (Days 14 and 56) captures early and intermediate dynamics but does not address the long-term stabilization of URT communities or resistome profiles. In addition, the low microbial biomass of nasal and oral samples increases susceptibility to contamination and technical noise, potentially affecting fine-scale taxonomic resolution. Finally, resistome quantification relied on relative abundance metrics rather than absolute gene copy numbers. Future studies integrating host immune profiling, metabolomics, and mechanistic modeling in larger cohorts will be essential to more fully elucidate the biological pathways underlying gut-lung microbial connectivity.

In summary, this longitudinal metagenomic analysis shows that FMT, beyond restoring gut microbial diversity and reducing intestinal antimicrobial resistance burden, is associated with microbial changes in the upper respiratory tract microbiome. The nasal niche exhibited transient reductions in richness and diversity, along with selective shifts in taxa commonly considered pathobionts and an associated decline in ARG abundance, whereas the oral microbiome remained ecologically stable. Together, these findings are consistent with microbial gut-lung connectivity and suggest that gut-targeted interventions may be associated with coordinated microbial and resistome changes across mucosal sites. Future studies incorporating extended follow-up, strain-resolved multi-omics approaches, and clinical respiratory outcomes will be essential to clarify the durability and functional significance of these associations.

## MATERIAL AND METHODS

### Study Design and Ethics

Human adult participants were enrolled in a longitudinal cohort at the University of Utah Hospital under the protocol titled “Profiling Changes in Respiratory Tract Microbiome in Intestinal Disease,” approved by the University of Utah IRB (IRB #00129810). The cohort consisted of fecal microbiota transplantation (FMT) recipients with recurrent *Clostridioides difficile* infection (CDI) and healthy community controls. Written informed consent was obtained from all participants. Patients were educated on sample collection techniques, and self-sampling occurred at Pre–FMT (Day 0) and Post–FMT on Days 14 and 56. Samples from healthy controls were collected at Day 0 and Day 14.

### Eligibility Criteria and Clinical Data

The inclusion criteria for FMT recipients were age over 18 years, ability to self-collect nasopharyngeal, oropharyngeal, and stool samples, and anticipated receipt of FMT for recurrent CDI, with permitted antibiotics (vancomycin PO, metronidazole, or fidaxomicin) administered within the prior 3 months. Individuals from either group were excluded if they had taken any antibiotics (other than those listed above) in the last month or were currently suffering from a respiratory tract infection. Individuals in the healthy control group were excluded if they had experienced gastrointestinal illness within the past month.

Demographics, comorbidities, lifestyle, and diet were self-reported and captured via a REDCap questionnaire (Table 1; Supplementary Table 1; **Supplementary Figure 1**).

### Sample Processing and DNA Extraction

Each fecal sample, nasal swab, and oral swabs were self-collected at each time point. Fecal samples were kept on ice, and oral and nasal swabs were collected into RNA later during transport to the laboratory. Upon arrival at the laboratory, samples were stored at -80 °C until processing. Fecal samples and swabs were vortexed for 3 minutes at high speed, and the remaining medium was centrifuged at 10,000 × g for 10 minutes to pellet microbial content. Microbial DNA was extracted using the ZymoBIOMICS™ 96 MagBead DNA Kit (Zymo Research, Irvine, CA, USA; Cat No. D4308), following the manufacturer’s protocol. Briefly, samples were transferred into lysis tubes containing BashingBeads and lysis buffer, followed by mechanical disruption using a bead beater (Fisherbrand™ Bead Mill 24 Homogenizer, Fisher Scientific, Waltham, MA, USA; Cat No. 15340163). Bead beating was performed in three cycles at 6 m/s for 1 minute each, with 5-minute resting intervals between cycles. Lysates were clarified by centrifugation, and DNA was bound to magnetic beads using the provided binding buffer. After sequential washing steps to remove contaminants, DNA was eluted in 50 µL of elution buffer. DNA concentration and purity were assessed using both a NanoDrop spectrophotometer and a Qubit fluorometer. Extracted DNA was stored at −80 °C until library preparation.

### Library Preparation and Sequencing

Library preparation and Whole-Genome Sequencing for 163 samples from 25 patients and donors were performed at the High-Throughput Genomics Core at the University of Utah. Metagenomic libraries were generated using the NEBNext® Ultra™ II FS DNA Library Prep Kit for Small Genomes (New England Biolabs, Ipswich, MA, USA), following the manufacturer’s instructions. This kit is optimized for whole-genome sequencing of low-input and fragmented DNA. Sequencing was conducted on the NovaSeq X Series platform using the 10B Reagent Kit, generating 150 × 150 bp paired-end reads. Each sample was sequenced to a target depth of at least 20 million reads.

### Processing sequencing data

Sequencing reads were demultiplexed using the GNomEx CORE Browser at the Huntsman Cancer Institute of the University of Utah. Trim Galore (v0.6.10) was used to trim adapters and filter low-quality reads. Reads were then mapped to the human genome (GRCh38) using BWA (v0.7.18). Reads that did not map to the human genome were then filtered with samtools (v1.1.6) and used for taxonomic analysis. In a separate pipeline for resistome characterization, host depletion was performed by aligning trimmed reads with Bowtie2 (v2.2.9). Unmapped reads were then aligned using the Resistance Gene Identifier (RGI, v6.0.5) to the Comprehensive Antibiotic Resistance Database (CARD, v4.0.1) using the KMA (v1.6.6) engine.

### Compositional, abundance, and resistome analysis

Taxonomic assignment was performed using Kraken2 (v2.1.2) against a custom database that combines the standard Kraken database with EuPathDB46 genomes. Kraken2 reports of inferred genus and species-level community composition were concatenated using Python (3.10.6). R (v4.4.3) was used to calculate Shannon’s diversity, Bray-Curtis dissimilarity between samples, dimensionality reduction with Principal Coordinate Analysis (PCoA), and differential abundance analysis. For all analyses, genus and species relative abundance were used, normalizing assigned read counts to the total number of assigned reads. For resistome analysis, gene mapping results from RGI were concatenated using Python and analyzed in R. R ggplot2 and pheatmap, as well as Graph Prism (v10.6.1), were used for plot generation. Code used for analysis is available upon request.

### Software and Reproducibility

Analyses were performed in R(v4.4.3) and Python (v3.10.6). Key packages: vegan (ordination/PERMANOVA), deseq2 (volcano plots) ggplot2, pheatmap (heatmaps).

### Statistical Analysis

Statistical Analyses were performed using R (v4.4.3), and Graph Prism (version 10.6.1). Figures were created with the ggplot2 and pheatmap libraries, as well as GraphPad Prism. Compositional analysis was performed by calculating Shannon’s diversity index, Pielou’s evenness, Abundance-based Coverage Estimator (ACE), and Bray-Curtis dissimilarity. PERMANOVA was used to determine the statistical significance of the PCoA plots. Sample groups were compared using a paired t-test with Benjamini-Hochberg p-value correction.

## DATA AVAILABILITY

The processed microbial data and diversity metrics are available upon request.

## Supporting information

Supplemental Material _Vallecillo et al

## ACKNOWLEDGEMENTS

The research was funded by the National Institutes of Health under Ruth L. Kirschstein National Research Service Award T32HL105321 (to MV-Z) and T32HG008962 (to DGB), and the Dr. Thomas D. Rees and Natalie B. Rees Presidential Endowed Chair in Global Medicine (to DTL). We would like to thank Zac Stephens, PhD (Department of Pathology, University of Utah), for bioinformatics assistance. We acknowledge support from the High-Throughput Genomics and Cancer Bioinformatics Shared Resource and HCI’s National Cancer Institute Cancer Center Support Grant P30CA042014. We also acknowledge partial support by the National Center for Advancing Translational Sciences of the National Institutes of Health under Award Numbers UL1 TR002538 and UM1 TR004409. Bioinformatics analysis was performed at the Utah Center for Genetic Discovery Core Facility, part of the Health Sciences Center Cores at the University of Utah. The support and resources from the Center for High Performance Computing at the University of Utah are gratefully acknowledged. The computational resources used were partially funded by the NIH Shared Instrumentation Grants S10OD034321 and S10OD021644. We also acknowledge and thank the study staff and the study participants who made this work possible.

## AUTHOR CONTRIBUTIONS

MLV-Z (Conceptualization, Data curation, Analysis, Investigation, Methodology, Visualization, Writing, the original draft, review, editing); DTL (Conceptualization, Funding, Project administration, Analysis, Writing, review, editing); AA (Software, Validation, Analysis, Visualization, Methodology, Writing, review, editing); DGB (Data curation, Analysis, Investigation, Methodology, Writing, review, editing); TAW (Data curation, Writing, review, editing); TH (Data curation, Writing, review, editing); MM (Software, Validation, Visualization, Writing, review, editing); KD (Data curation, Writing, review, editing); CN (Data curation, Writing, review, editing), AF (Methodology, Analysis, Writing, review, editing

## ETHICS DECLARATIONS

The authors declare no competing interests

## SUPPLEMENTAL INFORMATION

Supplemental Figure 1

Supplementary Table 1

