## Supplemental Material _Vallecillo et al for "Longitudinal Changes in Nasal and Oral Microbiome and Antimicrobial Resistance Gene Profiles in Response to Human Fecal Microbiota Transplantation"

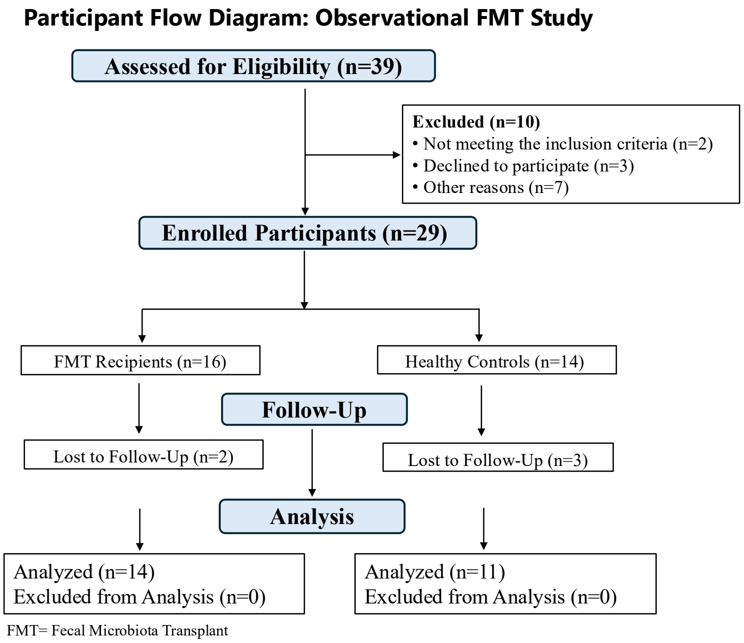

**Supplemental Fig. 1. Participant Flow Diagram for FMT Intervention Study**. Summary of participant progression through the study phases: screening, enrollment, group assignment, follow-up, and analysis.

**Supplementary Table 1. Lifestyle, supplement use, and dietary habits of FMT recipients and healthy controls.**

**A. Lifestyle Factors of FMT Recipients and Healthy Control Group**

| | **FMT Recipients** | **Healthy Control Group** |
|---|---|---|
| | **N=14 (56 %)** | **N=11 (44%)** |
| Alcohol use | 2 (14) | 6 (55) |
| Exercise habits | 10 (71) | 7 (64) |

**B. Supplement use of FMT Recipients and Healthy Control Group**

| | **FMT Recipients** | **Healthy Control Group** |
|---|---|---|
| | **n (%)** | **n (%)** |
| Multivitamin | 9 (64) | 4 (36) |
| Vit. D | 8 (57) | 3 (27) |
| Vit. B | 7 (50) | 2 (18) |
| Probiotics | 9 (64) | 3 (27) |

**C. Dietary Habits of FMT Recipients and Healthy Control Group**

| | **FMT Recipients** | **Healthy Control Group** |
|---|---|---|
| | **n = 14 (56%)** | **N=11 (44%)** |
| Vegetarian | 0 | 2 (18) |
| Meat eater | 13 (93) | 8 (72) |
| Poultry | 14 (100) | 8 (72) |
| Red meat | 13 (93) | 9 (82) |
| Seafood | 8 (57) | 5 (45) |
| Eggs | 14 (100) | 8 (72) |
| Dairy | 12 (86) | 9 (82) |
| Salted snacks | 13 (93) | 7 (64) |
| Whole grains | 13 (93) | 10 (91) |
| Vegetables | 13 (93) | 10 (91) |
| Sugary sweets | 13 (93) | 10 (91) |
| Sugar-sweetened beverages | 8 (57) | 5 (45) |
| Artificially sweetened beverages | 4 (29) | 5 (45) |
